# Proximal Exploration for Model-guided Protein Sequence Design

**DOI:** 10.1101/2022.04.12.487986

**Authors:** Zhizhou Ren, Jiahan Li, Fan Ding, Yuan Zhou, Jianzhu Ma, Jian Peng

## Abstract

Designing protein sequences with a particular biological function is a long-lasting challenge for protein engineering. Recent advances in machine-learning-guided approaches focus on building a surrogate sequence-function model to reduce the burden of expensive in-lab experiments. In this paper, we study the exploration mechanism of model-guided sequence design. We leverage a natural property of protein fitness landscape that a concise set of mutations upon the wild-type sequence are usually sufficient to enhance the desired function. By utilizing this property, we propose Proximal Exploration (PEX) algorithm that prioritizes the evolutionary search for high-fitness mutants with low mutation counts. In addition, we develop a specialized model architecture, called Mutation Factorization Network (MuFacNet), to predict low-order mutational effects, which further improves the sample efficiency of model-guided evolution. In experiments, we extensively evaluate our method on a suite of in-silico protein sequence design tasks and demonstrate substantial improvement over baseline algorithms.

## 1. Introduction

Protein engineering aims to discover novel proteins with useful biological functions, such as fluorescence intensity (Biswas et al., 2021), enzyme activity (Fox et al., 2007), and therapeutic efficiency (Lagassé et al., 2017), where the functions of a particular protein is determined by its amino-acid sequence (Crick, 1958; Nelson et al., 2008). The *protein fitness landscape* (Wright, 1932) characterizes the mapping between protein sequences and their functional levels (*aka*. fitness scores). It formulates the engineering of proteins as a sequence design problem. The evolution in nature can be regarded as a searching procedure on the protein fitness landscape (Smith, 1970), which is driven by natural selection pressure. This natural process inspires the innovation of *directed evolution* (Arnold, 1998), the most widely-applied approach for engineering protein sequences. It mimics the evolution cycle in a laboratory environment and conducts an iterative protocol. At each iteration, enormous variants are generated and scored by functional assays, in which sequences with high fitness are selected to form the next generation. Recent attempts focus on using machine learning approaches to build a surrogate model of protein landscape (Yang et al., 2019; Wittmann et al., 2021) and designing model-guided searching schemes (Biswas et al., 2018). This paradigm can effectively reduce the burden of laboratory experiments by performing in-silico population selection through model-based fitness prediction.

The exploration mechanism of directed evolution is a simple greedy strategy. It starts from the wild-type sequence and continuously accumulates beneficial mutations in a hill-climbing manner on the fitness landscape (Romero & Arnold, 2009). As the evolution is conducted in both the real landscape and the surrogate model, this unconstrained greedy search may lead to sequences far from the wild type. These sequences with high mutation counts are laborious to synthesize for mass production (Storici & Resnick, 2006; Fowler et al., 2014). In addition, it is hard for the training procedure of the surrogate model to reuse previous samples that are far from the current search region.

However, as shown in Figure 1, the protein landscape is known to be sparse in the vast sequence space, where the local landscape near the wild-type point is rugged and multi-peaked. It implies that a few amino-acid mutations on key positions are sufficient to dramatically alter the fitness score of the protein (Weinreich et al., 2005). Searching through regions with low mutations around wild-type sequence can improve searching effectiveness and reduce wet-lab labor.

**Figure 1.**
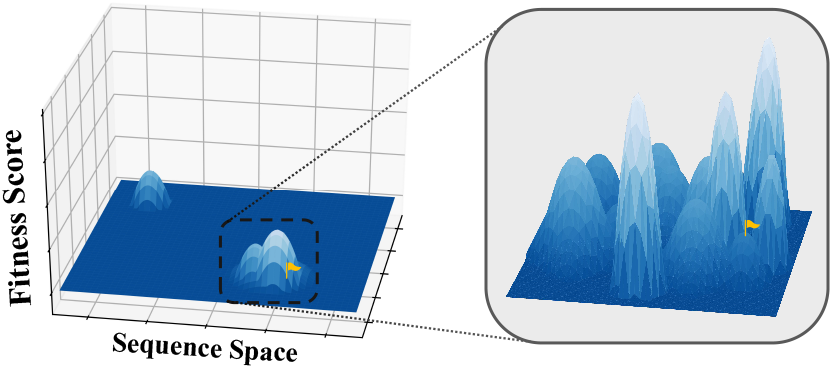
(Left) Functional proteins are rare in the sequence space. The yellow flag corresponds to the wild-type sequence. (Right) The local landscape near the wild type is rugged. A few amino-acid mutations may dramatically alter the protein fitness.

Regarding the natural property of the protein landscape, we propose a novel exploration mechanism that goes beyond the classical paradigm of directed evolution. Our exploration method, named *Proximal Exploration* (PEX), aims to search for effective candidates of low-order mutants (*i*.*e*., mutants near the wild-type sequence) rather than greedily accumulate mutations along the evolutionary pathway. By formulating the local search around the wild-type sequence as a proximal optimization problem (Parikh & Boyd, 2014), we derive a regularized objective for searching the steepest improvement upon the wild type with the limited counts of amino-acid mutations. Furthermore, we design a specialized network architecture, called *Mutation Factorization Network* (MuFacNet), to model low-order mutational effects based on local neighbor information, which additionally improves the sample efficiency of our model-guided sequence design. Our method shows great performance over baselines on multiple protein sequence design benchmarks.

## 2. Related Work

### ML for Protein Landscape Modeling

Advances in experimental technologies, such as deep mutational scanning (Fowler & Fields, 2014), yield large-scale generation of mutant data. It enables data-driven approaches to model the protein fitness landscape, which is one of the crucial problems for protein engineering. Recent works start to focus on leveraging co-evolution information from multiple sequence alignments to predict fitness scores (Hopf et al., 2017; Riesselman et al., 2018; Luo et al., 2021) and utilizing pre-trained language models to conduct transfer learning or zero-shot inference (Rao et al., 2019; Alley et al., 2019; Meier et al., 2021; Hsu et al., 2022). The learned ML model can be used to screen enormous designed sequences in silico (Ogden et al., 2019; Bryant et al., 2021; Gruver et al., 2021; Shan et al., 2022), which serves as a surrogate of expensive wet-lab experiments.

### Exploration Algorithms for Sequence Design

Directed evolution is a classical paradigm for protein sequence design, which has achieved several successes (Bloom & Arnold, 2009; Dalby, 2011; Arnold, 2018). Under this framework, machine learning algorithms play an important role in improving the sample efficiency of evolutionary search. Angermueller et al. (2020a) proposes an ensemble approach that performs online adaptation among a heterogeneous population of optimization algorithms. Angermueller et al. (2020b) formulates sequence design as a sequential decision-making problem and conducts model-based reinforcement learning for efficient exploration. Brookes & Listgarten (2018) and Brookes et al. (2019) use adaptive sampling to generate high-quality mutants for batch Bayesian optimization. Zhang et al. (2021) proposes a probabilistic framework that unifies black-box sequence design and likelihood-free inference and presents a methodology to develop new algorithms for sequence design.

### Proximal Optimization

Proximal methods are a family of optimization algorithms that decompose non-differentiable large-scale problems to a series of localized smooth optimization (Parikh & Boyd, 2014). The most representative approach is proximal gradient (Combettes & Wajs, 2005), which conducts a convex step-wise regularization to smooth the optimization landscape without losing the optimality of convergence. It is related to natural gradient (Amari, 1998) and trust-region methods (Sorensen, 1982). These methods aim to find the steepest improvement direction in a restricted small region. In this paper, we adopt the formulation of proximal methods to deploy localized optimization around the wild-type sequence. We propose an exploration algorithm to find proximal maximization points (Rockafellar, 1976) for black-box sequence design.

## 3. Problem Formulation and Background

### Protein Sequence Design upon Wild Type

The protein sequence design problem is to search for a sequence *s* with desired property in the sequence space 𝒱^*L*^, where 𝒱 denotes the vocabulary of amino acids and *L* denotes the desired sequence length. We aim to maximize a protein fitness function *f* : 𝒱^*L*^ → ℝ by editing sequences. The protein fitness mapping *f*(*s*) is a black-box function that can only be measured through wet-lab experiments. As a reference to the design problem, a wild-type sequence *s*^wt^ is given as the starting point of sequence search, which is the sequence occurring in nature. In general, the wild type *s*^wt^ has already expressed decent fitness under the natural selection evolutionary process towards the target function (Fisher, 1958; Chari & Dworkin, 2013). In this paper, we focus on the modification upon the wild-type sequence *s*^wt^ to enhance the desired protein function. Specifically, we aim to design low-order mutants that have low mutation counts compared to the wild-type sequence.

### Batch Black-Box Optimization

Protein sequence design with respect to a fitness function *f*(*s*) is a batch black-box optimization problem. Advances in experiment technology have enabled the parallel fitness measurement over a large batch of sequences (Kosuri et al., 2013; Peterman & Levine, 2016). However, each round of parallel experiments in the wet lab is expensive and time-consuming. It leads to an online learning problem with high throughput but limited interactions. Formally, we conduct *T* rounds of batch optimization with batch size *M* ≫ *T* for sequence-fitness measurements within each round.

### Model-directed Evolution

Building a surrogate model for in-silico evolutionary selection is an effective approach to reduce the burden of expensive wet-lab experiments. It trains a fitness model 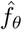, parameterized by *θ*, to predict the fitness of mutant sequences. More specifically, the surrogate model optimizes the following regression loss for fitness prediction:

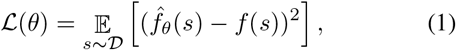

where 𝒟 is a dataset of experimentally measured sequences. The learned surrogate model 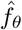 can be used to predict the fitness of unseen sequences, which can thus guide in-silico sequence searching and improve the sample-efficiency of directed evolution.

## 4. Proximal Exploration for Sequence Design

In this section, we introduce our approach, Proximal Exploration (PEX), which prioritizes the search for high-fitness sequences with low mutation counts upon the wild type. We formulate this localized search as a proximal optimization problem and implement the exploration mechanism through a model-guided approach. In addition, we propose a specialized network architecture to model the local landscape around the wild type, which further improves the efficiency of proximal exploration.

### 4.1. Overview of Proximal Exploration

In the natural evolutionary process, it is usually sufficient for a protein to significantly enhance its fitness score by mutating only a few amino acids within a long sequence. Motivated by this natural principle, we consider restricting the search space near the wild type and seek for high-fitness mutants with low mutation counts. Based on the paradigm of directed evolution, the basic idea is to adopt a regularized objective that prioritizes low-order mutants in the sequence selection step. *i*.*e*., our algorithm prefers to select sequences with low mutation counts for artificial evolution. Formally, we employ the proximal optimization framework to formulate such a localized searching scheme.

#### Evolution via Proximal Optimization

Following the formulation of proximal optimization (Parikh & Boyd, 2014), we define a regularized objective function that penalizes the deviation from the wild-type sequence *s*^wt^:

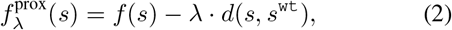

where *f*(·) is the original objective function, *i*.*e*., the protein fitness score. λ ≥ 0 denotes the regularization coefficient. *d*(·, ·) denotes a distance metric in the sequence space. In this paper, without further specification, the metric *d*(·, ·) refers to evolutionary distance (*i*.*e*., Levenshtein edit distance). We aim to find a *proximal maximizer* (or *proximal point* (Rockafellar, 1976)) with respect to the wild-type sequence *s*^wt^:

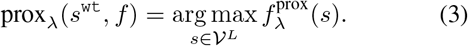

The value of λ controls the weight of proximal regularization. With larger values of λ, the searching procedure driven by Eq. (2) would become more concentrated around the wild-type sequence, and go far otherwise. The degraded case with λ = 0 is equivalent to the maximization of the original objective function: prox_λ=0_(*s*^wt^, *f*) = arg max_*s*∈𝒱_^*L*^ *f*(*s*). By varying the value of regularization coefficient λ, we define the *proximal frontier* as follows:

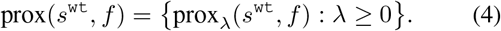

#### Geometric Interpretation

Figure 2a shows a geometric illustration of proximal frontier prox(*s*^wt^, *f*). It interprets why this set of proximal points is named as a *frontier*. We plot a coordinate system where each protein sequence is represented by its fitness score *f* (*s*) and its distance *d*(*s, s*^wt^) to the wild-type sequence *s*^wt^. The sequences in prox(*s*^wt^, *f*) are located at the upper frontier on the convex closure of valid sequence space. These proximal points are Pareto-efficient solutions to trade off between maximizing protein fitness score *f* (*s*) and minimizing the evolutionary distance *d*(*s, s*^wt^) to the wild-type sequence, *i*.*e*., the trade-off between the major objective and the regularization term. The sequences lying on the left-upper region of this coordinate system correspond to the mutants with high fitness and low mutation counts, which is the goal of our method.

**Figure 2.**
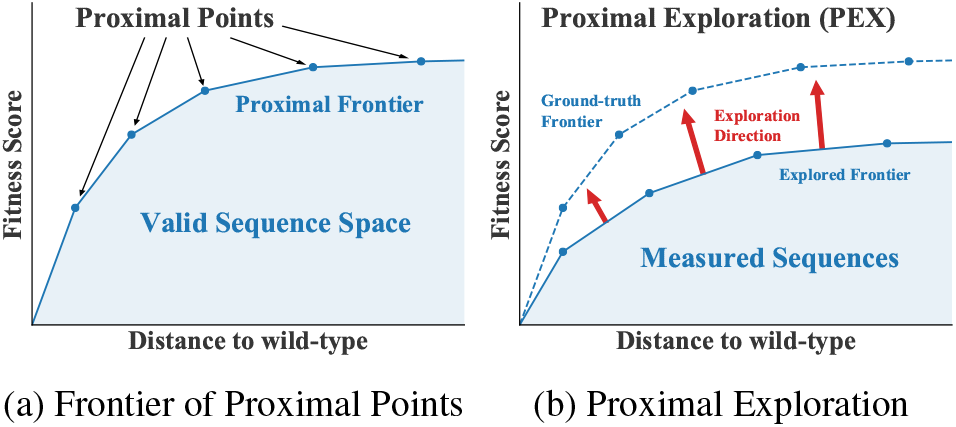
(a) A geometric illustration of the definition of proximal points and proximal frontier. (b) The goal of proximal exploration is finding the ground-truth proximal frontier.

#### Proximal Exploration (PEX)

In classical evolutionary algorithms, only top-fitness sequences are selected and mutated within the evolution cycle. Such a greedy evolutionary process hinders the efficiency of exploration and does not utilize the property of the natural protein fitness landscape. We consider a different mechanism that performs exploration along the proximal frontier and seeks high-fitness mutant sequences with low mutation counts. As shown in Figure 2b, our exploration algorithm, Proximal Exploration (PEX), aims to extend the proximal frontier instead of only optimizing the fitness score. We consider this proximal exploration mechanism as a regularization of the search space of protein sequences.

##### Algorithm 1

Proximal Exploration (PEX)

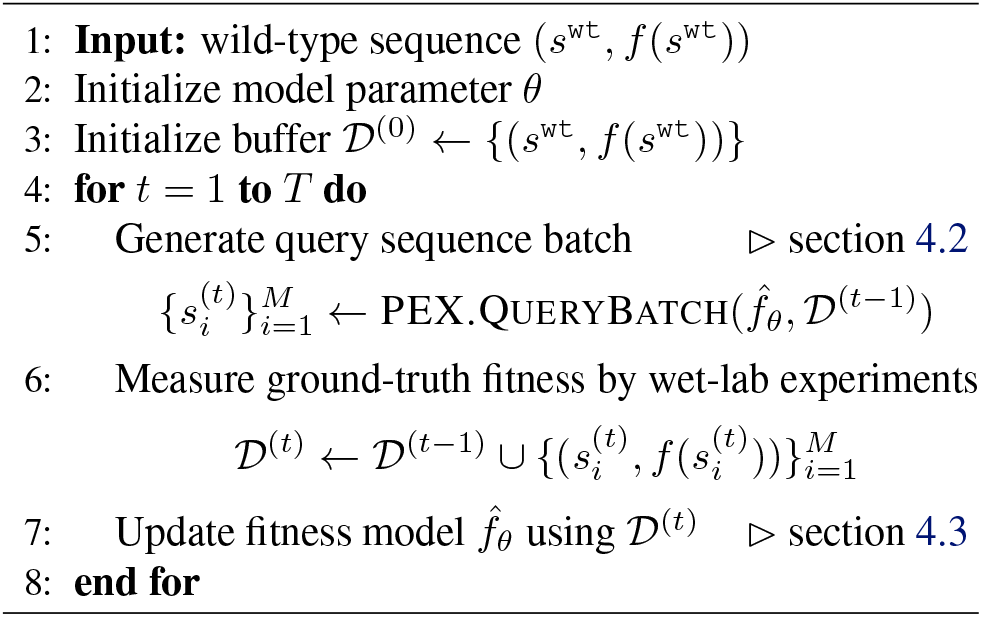

The overall procedure of our algorithm is summarized in Algorithm 1. Following the problem formulation introduced in section 3, our algorithm conducts *T* rounds of interaction with the laboratory. At each round, we propose a query batch containing *M* sequence candidates (line 5) and measure their fitness scores through the wet-lab experiments (line 6). The measured protein fitness data are used to refine the fitness model 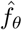 where *θ* denotes the model parameter (line 7). The major technical improvement of our method contains two parts:

1. We formalize the mechanism of proximal exploration through a model-guided approach (section 4.2).
2. We design a specialized model architecture to predict low-order mutational effects of proximal points, which further boosts the efficiency of model-guided exploration (section 4.3).

### 4.2. Model-guided Exploration on Proximal Frontier

In each round of batch black-box optimization, the exploration algorithm is required to generate a query batch given the experimental measurements collected in previous rounds. More specifically, a model-guided exploration algorithm constructs the query batch based on the fitness model 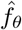 and a dataset of measured sequences 𝒟. The query generation procedure of PEX is summarized in Algorithm 2. The same as other model-guided methods, we first mutate sequences in measured dataset 𝒟 to obtain a large set of random sequences 𝒟^pool^ (*i*.*e*., a candidate pool of offspring in evolution) and then perform a model-based selection over these mutants.

#### Algorithm 2

PEX.QueryBatch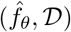

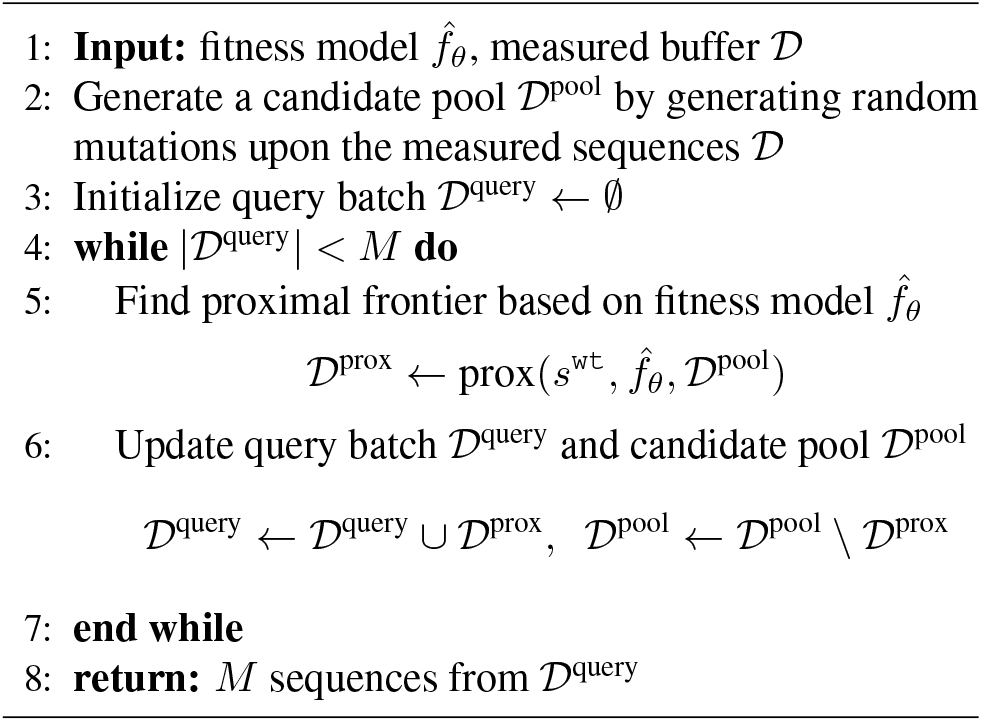

Classical evolutionary algorithms perform a greedy selection to determine the query batch. The selection criteria is given by the model fitness prediction 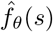. *i*.*e*., only sequences with high predicted fitness scores will be sent to laboratory measurements. In comparison, PEX considers a different objective to construct the query batch. To expand the frontier shown in Figure 2b, we compute the proximal frontier within the candidate pool 𝒟^pool^:

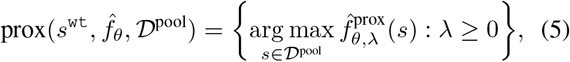

which is expected to explore Pareto-efficient sequences with either higher fitness or lower mutation count.

At each round of interaction with the wet lab, the query budget can support the experimental measurement of *M* sequences. In addition to the sequences exactly lying on the proximal frontier defined in Eq. (5), we diversify the query batch by including the nearby region of the frontier. We conduct an iterative query generation mechanism as shown in Algorithm 2. We iteratively compute the proximal frontier over the current candidate pool 𝒟^pool^ (line 5) and remove them from the pool (line 6). It enables us to effectively deploy proximal exploration in the batch-query setting.

#### Remark

The computation of 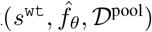 defined in Eq. (5) is equivalent to find the upper convex closure of 𝒟^pool^ in the coordinate system illustrated by Figure 2a, where the fitness scores are predicted by the surrogate model 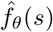. In our implementation, we use Andrew’s monotone chain convex hull algorithm (Andrew, 1979), which can be completed in *O*(*n* log *n*) time with *n* = | 𝒟^pool^|. Implementation details are included in Appendix A.2.

### 4.3. Modeling the Landscape of Low-order Mutants

One challenge of landscape modeling is the high sensitivity of protein fitness concerning amino-acid mutations. As illustrated in Figure 1, a few amino-acid substitutions may dramatically alter the fitness scores. Classical model architectures, such as CNN and RNN, represent the protein landscape as a sequence-fitness mapping. Since the fitness function is highly non-smooth in the sequence space, these models usually require a large amount of mutant data to capture the complicated mutational effects, which do not meet the demand of sequence design in data efficiency.

To address the challenge of non-smoothness for landscape modeling, we consider to leverage the algorithmic property of proximal exploration. Note that our exploration mechanism prioritizes the search for low-order mutants close to the wild-type sequence. This special property motivates us to design a specialized network architecture to model the local fitness landscape around the wild type. We propose a Mutation Factorization Network (MuFacNet) that factorizes the composition of mutational effects to the interactions among single amino-acid mutations. The major characteristic of MuFacNet is its input representation. It represents a mutant sequence by the corresponding set of amino-acid mutations upon the wild type, which emphasizes the effects of each single mutation site.

#### Forward View: Aggregation

Figure 3 presents the architecture of MuFacNet. Given an input mutant sequence, we first locate the sites of mutations in comparison to the wild type. We take the neighbor fragment of each mutated amino acid as its context information, *i*.*e*., a context window centered at the mutated amino acid. Each context window is then fed into an encoder network to generate the feature vector of the corresponding mutation site. We would obtain *K* separate feature vectors for *K* mutations, which are computed by a shared encoder network. These feature vectors encode the functionality of the given mutations. We perform a sum-pooling operator to aggregate them into a unified vector that represents the joint effect of mutations. Finally, the protein fitness prediction is given by a decoder network taking the joint-effect vector as its input.

**Figure 3.**
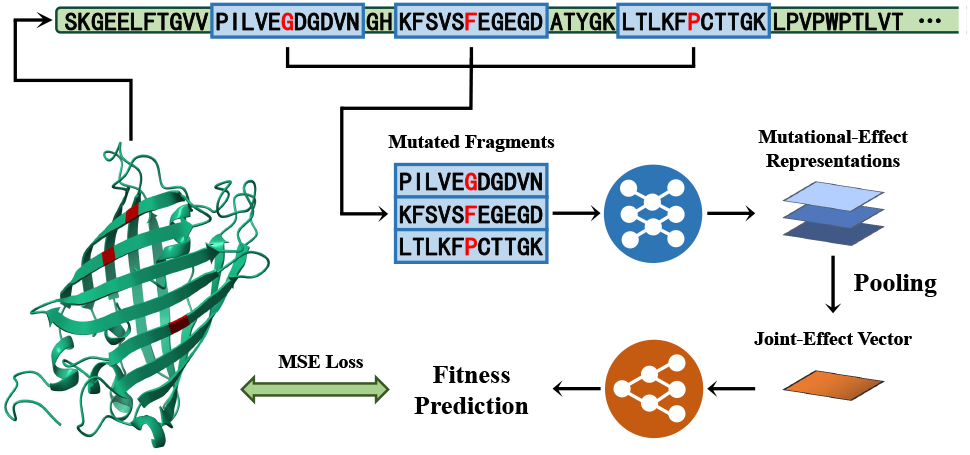
The model architecture and the forward flow graph of Mutation Factorization Network. The red marked sites are mutated amino acids upon the wild type. The illustrative example corresponds to the green fluorescent protein (PDB code: 2O29).

#### Backward View: Factorization

The supervision of Mu-FacNet is the same as the standard model training procedure. It learns from sequence-fitness data through an end-to-end paradigm. The data obtained in wet-lab measurements only indicate the joint effects of mutation compositions, and the design purpose of MuFacNet is to factorize the joint fitness value to local representations of every single mutation site. The pooling operator is the core component to realize this joint-to-local factorization. It characterizes the interaction among mutation sites through the arithmetical composition of their feature vectors.

#### Remark

The architecture of MuFacNet is a variant of set model that represents functions defined on sets. The sum-pooling operator ensures the model output is invariant to the order of input mutation context windows. Zaheer et al. (2017) proves that such a sum-pooling aggregation operator can preserve the universal approximability of neural networks. *i*.*e*., with sufficiently expressive encoder and decoder, a set model with sum-pooling aggregation can represent any functions mapping sets to values. This property ensures that MuFacNet can learn non-additive epistasis effects.

## 5. Experiments

In this section, we present experiments to evaluate the performance of our method. We show that PEX together with MuFacNet achieves state-of-the-art performance and significantly outperforms baselines in section 5.2 and 5.3. In addition, we conduct several ablation studies to investigate the functionality of each designed component, including the design principle of MuFacNet and the local-search mechanism of PEX.

### 5.1. Protein Engineering Benchmarks

We evaluate our exploration algorithm and baseline methods on a suite of in-silico protein engineering benchmarks. Following prior works, we simulate the ground-truth protein fitness landscape by an oracle model in replace of the wet-lab measurements. As the black-box optimization setting defined in section 3, the exploration protocol can only interact with the simulated landscape through batch queries. The sequence design algorithm cannot obtain any extra information about the black-box oracle.

We collect several large-scale datasets from previous experimental studies of protein landscape and use TAPE (Rao et al., 2019) as an oracle model to simulate the landscape. These fitness datasets involve extensive applications of protein engineering, including biosensors design, thermostability improvement, and industrial enzyme renovation. It enables us to formally compare the performance of exploration algorithms in an in-silico sandbox. The source code of our algorithm implementation and oracle landscape simulation models are available at https://github.com/ HeliXonProtein/proximal-exploration.

#### Green Fluorescent Proteins (avGFP)

Green Fluorescent Proteins from *Aequorea victoria*, which can exhibit bright green fluorescence when exposed to light in the blue to the ultraviolet range, are frequently used as biosensors to detect gene expressions and detect protein locations. Sarkisyan et al. (2016) assayed the fluorescence levels of nearly 52000 genotypes of avGFP obtained by random mutagenesis to the 239-length wild-type sequence, used as our landscape. Our goal is to optimize sequences with higher log-fluorescence intensity values in this 20^238^ search space.

#### Adeno-associated Viruses (AAV)

Engineering a 28-amino acid segment (position 561-588) of the VP1 protein located in the capsid of the Adeno-associated virus has drawn lots of attention in ML-guided design, aiming to remain viable for packaging of a DNA payload for gene therapy (Ogden et al., 2019). We adopt a library with approximately 284, 000 mutant data developed by Bryant et al. (2021) to build the landscape oracle. We aim to design more capable sequences as gene delivery vectors measured by AAV viabilities. The size of search space is 20^28^.

#### TEM-1 *β*-Lactamase (TEM)

TEM-1 *β*-Lactamase protein resisting to penicillin antibiotics in *E*.*coli* is widely studied to understand mutational effect and fitness landscape (Bershtein et al., 2006; Jacquier et al., 2013). We collect the fitness data from Firnberg et al. (2014) and Gonzalez & Ostermeier (2019), resulting in 17857 sequences as the landscape training data, where fitness is the view as thermodynamic stability. The optimization problem is to propose high thermodynamic-stable sequences upon wild-type TEM1 in the search space with size 20^286^.

#### Ubiquitination Factor Ube4b (E4B)

Ubiquitination factor Ube4b plays an important role in the trash degradation process in the cell by interacting with ubiquitin and other proteins. Starita et al. (2013) scored approximately 100, 000 mutation protein sequences by measuring their rates of ubiquitination to the target protein. We focus on designing E4B with higher enzyme activity on the landscape above. The size of search space is 20^102^.

#### Aliphatic Amide Hydrolase (AMIE)

Amidase encoded by *amiE* is an industrially-relevant enzyme from *from Pseudomonas aeruginosa*. By quantifying the growth rate of each bacterial strain carrying specific amidase mutants, Wrenbeck et al. (2017) measures 6629 sequences with single mutations to model the fitness landscape. We seek for optimizing amidase sequences which lead to great enzyme activities. It defines a search space with 20^341^ sequences.

#### Levoglucosan Kinase (LGK)

Levoglucosan kinase converts LG to the glycolytic intermediate glucose-6-phosphate in an ATP-dependent reaction. A mutant library containing 7891 mutated protein has been established and evaluated using the deep mutational scanning method to recognize mutational effects (Klesmith et al., 2015). Optimizing such protein for improved enzyme activity fitness is our goal. The size of search space is 20^439^.

#### Poly(A)-binding Protein (Pab1)

Some proteins function by binding to other biomolecules, *e*.*g*., RNA or DNA. Pab1 is one of them that binds to multiple adenosine monophosphates (poly-A) using the RNA recognition motif (RRM). Melamed et al. (2013) develop high throughput screening to assay the binding fitness of around 36000 double mutants of Pab1 variants in RRM region. As above, we desire to improve the binding fitness by introducing beneficial mutations to the wild-type sequence. The search space size is 20^75^ on a segment of the wild-type sequence.

#### SUMO E2 conjugase (UBE2I)

Utilizing variants to function mapping of human genomes is substantial for scientific research and clinic treatment. We put to use around 2000 variants of the disease-relevant protein, human SUMO E2 conjugase constructed by Weile et al. (2017) and boost the fitness measured by growth rescue rate at high temperature in a yeast strain. The size of search space is 20^159^.

### 5.2. Performance Comparison to Baseline Algorithms

We compare the performance of PEX with several baseline algorithms to design functional proteins by exploring the fitness landscape.

#### AdaLead

(Sinai et al., 2020) is an advanced implementation of model-guided evolution. At each round of batch query, AdaLead first performs a hill-climbing search on the learned landscape model and then queries the sequences with high predicted fitness. Such a simple implementation of model-directed evolution has been demonstrated to achieve competitive performance against many elaborate algorithms.

#### DyNA PPO

(Angermueller et al., 2020b) formulates protein sequence design as a sequential decision-making problem and uses model-based reinforcement learning to perform sequence generation. It applies proximal policy optimization (PPO; Schulman et al., 2017) to search sequences on a learned landscape model. Different from our proximal approach, PPO performs a regularization between the policies of two consecutive rounds. Since a slight alter in policy space may dramatically change the sequence generation distribution, DyNA PPO does not restrict the search space around wild type.

#### DbAS

(Brookes & Listgarten, 2018) establishes a probabilistic framework that uses model-based adaptive sampling to explore the fitness landscape. It trains a variational auto-encoder (VAE; Kingma & Welling, 2013) to model the distribution of high-fitness sequences and performs model-guided evolution using this sequence sampler.

#### CbAS

(Brookes et al., 2019) follows the same formulation as DbAS and consider a regularization to stabilize the model-guided search. The motivation and the regularization objective of CbAS are similar to the consideration of DyNA PPO. It restricts the sampling distribution to be supported by the given labeled dataset, which leads to a trust-region search concerning the learned surrogate model. It prevents the exploration algorithm from being trapped in the ill-posed regions with poor model generalization.

#### CMA-ES

(Hansen & Ostermeier, 2001) is a famous algorithm for evolutionary search. It is a second-order approach that estimates the covariance matrix to adaptively adjust the search strategy of the upcoming generations.

#### Bayesian Optimization

(BO; Mockus, 2012) is a classical paradigm for the sequential design problem. We adopt the implementation developed by Sinai et al. (2020), which uses an ensemble of models to estimate the uncertainty and construct the acquisition function for exploration.

We consider an ensemble of three CNN models as the default configuration for these model-guided approaches. Specifically, following the implementation of Angermueller et al. (2020b), DyNA PPO uses a hybrid ensemble and conducts an adaptive selection to pick reliable model candidates. In the evaluation of proximal exploration, **PEX+MuFacNet** and **PEX** refer to using an ensemble of three MuFacNets and the default CNN models, respectively. An ablation study on the model configuration is presented in section 5.3.

#### Performance Comparison

The protein engineering bench-marks introduced in section 5.1 have much longer sequence length than the proteins considered by previous works, which raises a challenge since the size of search space is exponential to the sequence length. The experiment results are presented in Figure 4. It shows that PEX generally outperforms baseline algorithms with significant improvement. Our approach can find sequences with higher fitness scores using fewer rounds of black-box queries, which demonstrates the improvement of sample efficiency. In Appendix B, we conduct an additional suite of performance evaluation with oracle landscape simulators based on ESM-1b (Rives et al., 2021), and our method also significantly outperforms baselines.

**Figure 4.**
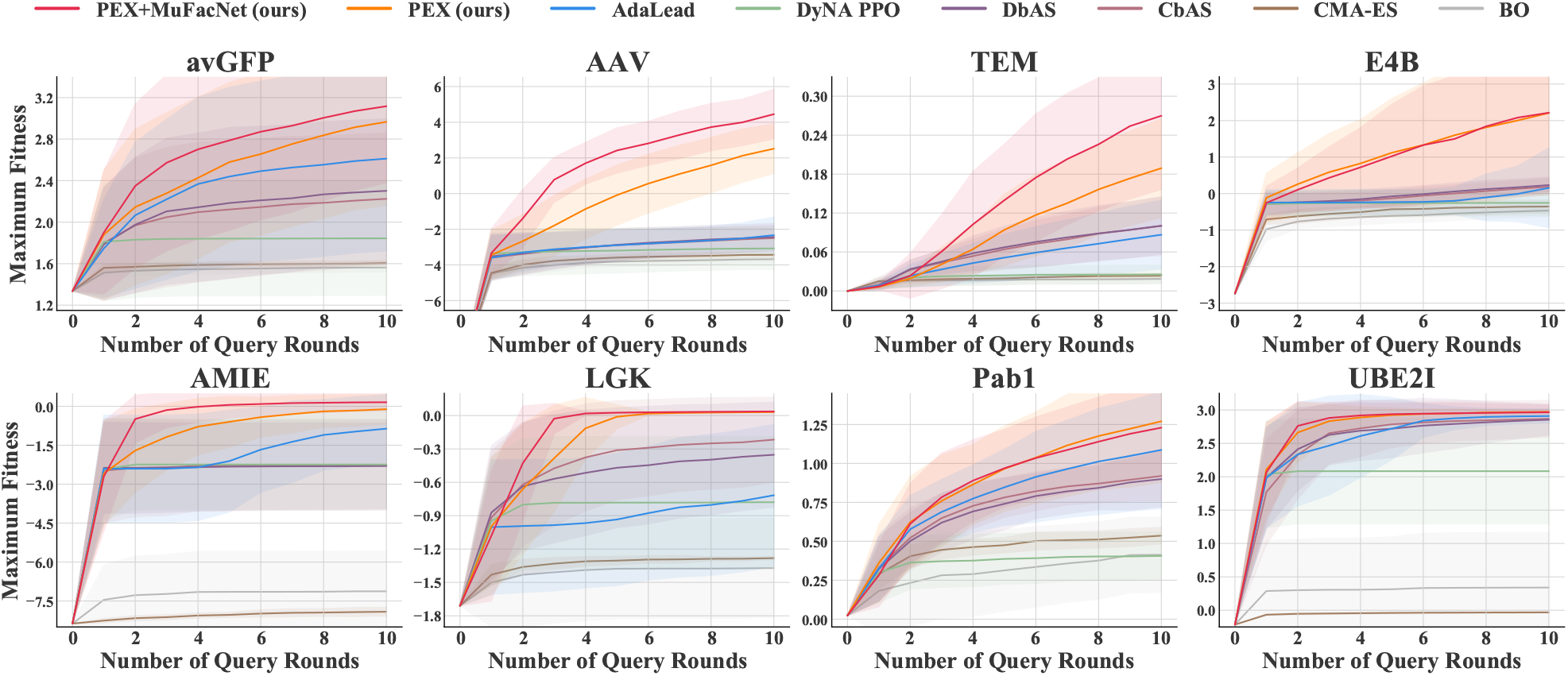
Learning curves on a suite of protein engineering benchmark tasks. Each round of batch black-box query can measure the fitness scores of 100 sequences. The evaluation metric is the cumulative maximum fitness score among queried sequences. All curves are plotted from 80 runs with random network initialization. The shaded region indicates the standard deviation.

### 5.3. Effectiveness of MuFacNet

The performance of a model-guided algorithm depends on the quality of its back-end surrogate model. To establish an ablation study on the model configuration, we consider three types of model architectures: CNN, RNN, and MuFacNet, where all of these models are implemented by an ensemble of three network instances. CNN with 1D-convolution is a popular choice for protein fitness prediction (Shanehsaz-zadeh et al., 2020), and we adopt the implementation developed by AdaLead (Sinai et al., 2020). The RNN architecture is adopted from Gruver et al. (2021), which uses stacked GRUs (Chung et al., 2014). The implementation details of MuFacNet are included in Appendix A.3.

The experiment results regarding different model architectures are presented in Table 1. The **Overall** evaluation refers to the normalized scores averaged overall tasks. *i*.*e*., we normalize the cumulative maximum fitness score of each task to the unit interval [0, 1] and present the average overall tasks. The results show that **PEX+MuFacNet** provides the most stable performance. It achieves the best average performance in 6 out of 8 benchmark tasks and only marginally underperforms in the remaining 2 tasks. MuFacNet can also improve the performance of AdaLead, DbAS, and CbAS. It shows that MuFacNet is a widely applicable model for sequence design based on the given wild type.

**Table 1.**
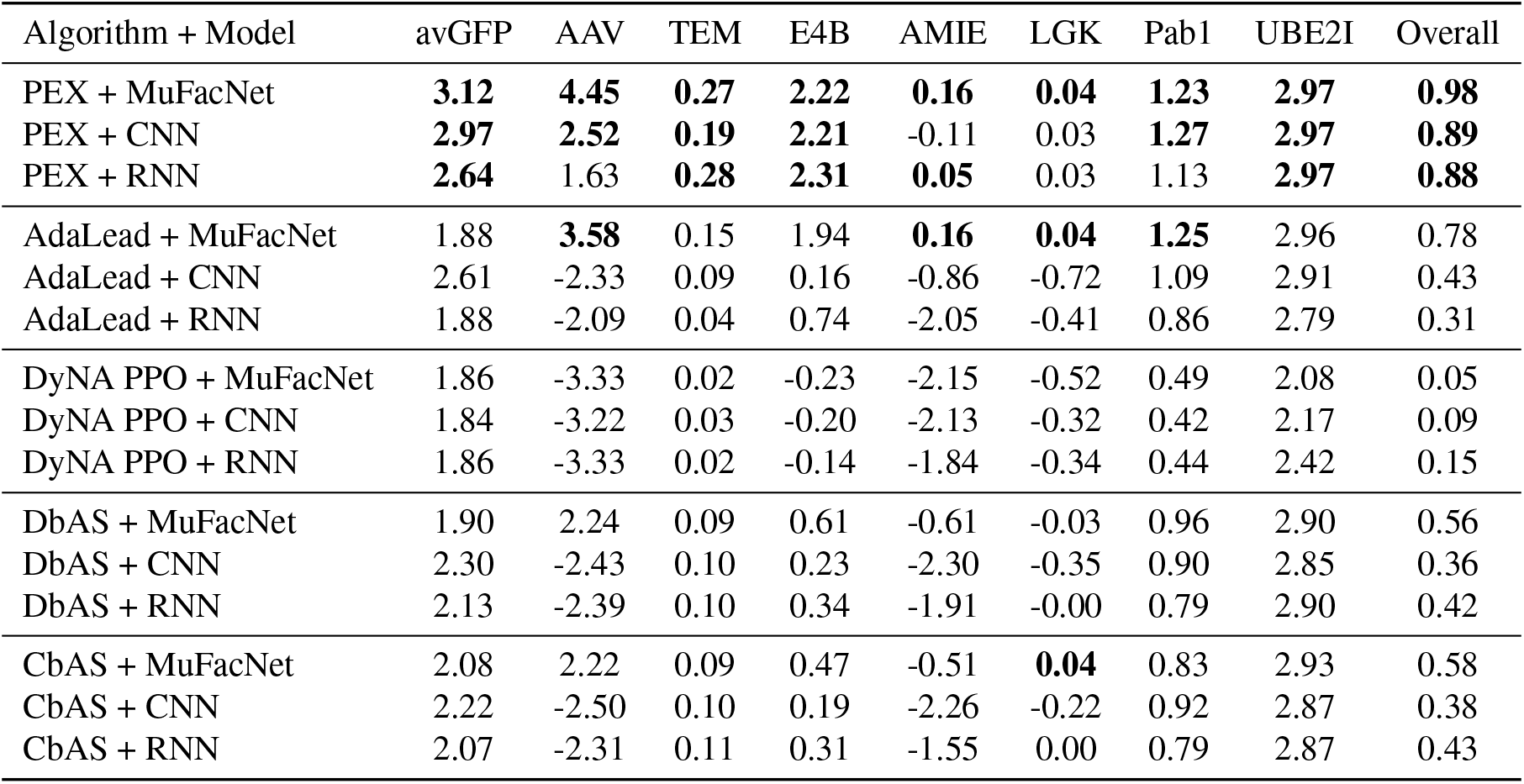
Comparing different model architectures for model-guided sequence design. We present the **maximum fitness scores** obtained in 10 rounds of black-box queries. The **top-3 algorithms** of each task are marked by the bold font. “Overall” refers to the normalized score averaged over 8 tasks. Every entry of this table is averaged over 80 runs. The learning curves are included in Appendix C.

### 5.4. Encoding Context Information of Mutation Sites

The major characteristic of MuFacNet is its input representation that represents a mutant sequence by a set of amino-acid mutations upon the wild type. To demonstrate the superior performance of MuFacNet is supported by such a set-representation, we conduct several ablation studies to investigate the module design of MuFacNet. First, we vary the size of the context window to locate the mutation site. In the default configuration, we take context windows with radius 10 centered at each mutation site as the inputs of MufacNet. In addition, we consider an alternative way to represent the mutated amino acids. We concatenate the positional encoding (Vaswani et al., 2017) with size 32 to the one-hot encoding of the amino-acid mutation and use an MLP-encoder to produce the feature vector of mutation sites. The experiment results are presented in Table 2. The typical implementation of mutation encoding does not largely affect the performance of the downstream protein sequence design. It indicates that the paradigm of set-representation is critical to the performance improvement. In this paper, since we focus on learning from a small set of protein-fitness data, the encoder and decoder architectures of MuFacNet are implemented by simple modules such as 1D-CNN and MLP layers. A future work is to investigate advanced set models (Lee et al., 2019) on large-scale protein fitness datasets.

**Table 2.** Ablation studies on the methods to represent the context information of each single amino-acid mutation site. The learning curves of all eight landscapes are included in Appendix D.

| Mutation Encoding | avGFP | AAV | TEM | E4B |
| --- | --- | --- | --- | --- |
| Window (radius=5) | 2.96 | 4.68 | 0.23 | 2.62 |
| Window (radius=10) | 3.12 | 4.45 | 0.27 | 2.22 |
| Window (radius=15) | 2.86 | 4.58 | 0.25 | 2.40 |
| Positional Encoding | 3.15 | 4.32 | 0.27 | 2.61 |
| without MuFacNet | 2.97 | 2.52 | 0.19 | 2.21 |

### 5.5. Discovering Low-Order Mutants with High Fitness

In addition to the ability to discover high-fitness proteins, we present another strength of PEX, *i*.*e*., the sequences designed by our method have a low mutation count upon the wild-type sequence. To demonstrate this algorithmic property, we collect the best sequence designed by every algorithm and evaluate their mutation counts, *i*.*e*., the number of amino-acid mutations required to synthesize these sequences from the wild type sequence. The results are presented in Figure 5. In comparison to baseline algorithms, PEX can discover high-fitness sequences with far less number of mutations upon the wild type. Since synthesizing sequences with a large number of mutation sites is time- and labor-consuming for modern site-directed mutagenesis toolkit (Hogrefe et al., 2002; Carrigan et al., 2011), and mutants with fewer mutation count may also have good performance in other biochemical aspects (Klesmith et al., 2017; Araya & Fowler, 2011), our framework is a more powerful tool for efficient high-throughput protein design.

**Figure 5.**
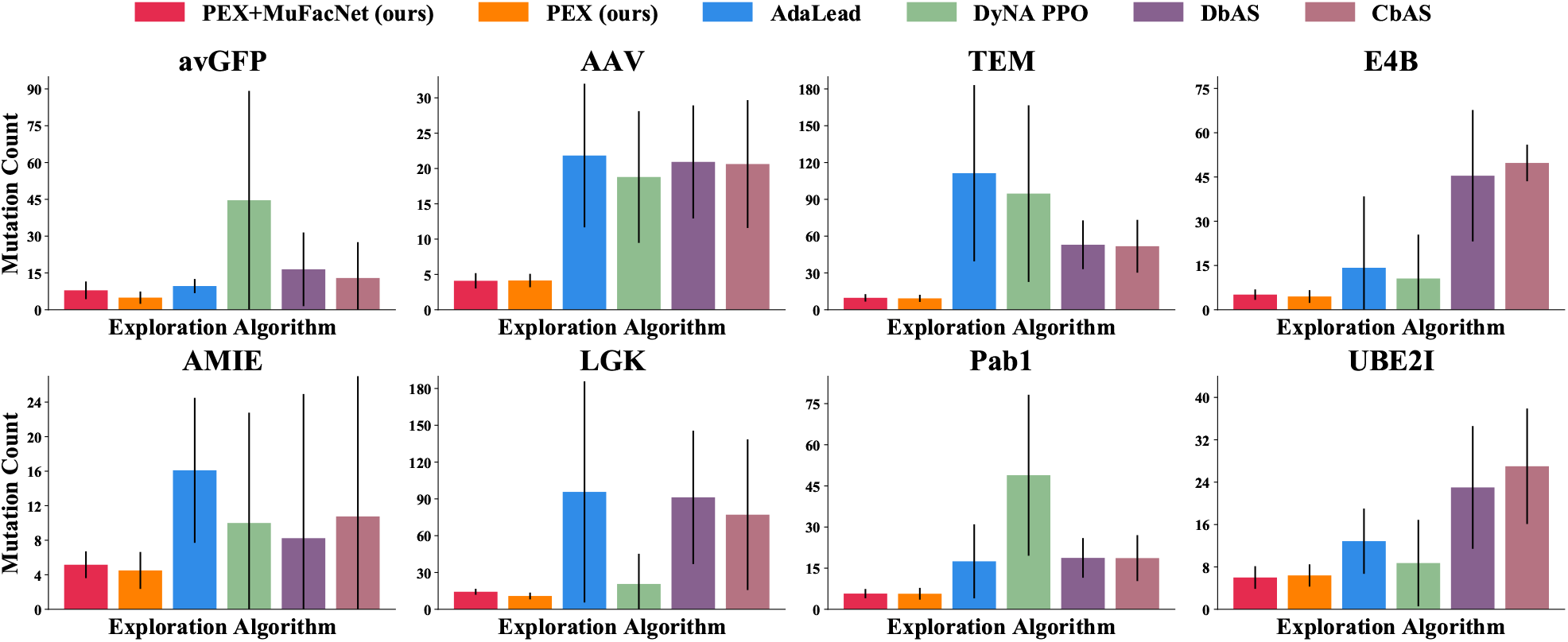
The mutation count of the best-designed sequence for each method in multiple benchmarks. Our methods achieve fewer mutation count while maintaining high fitness compared to other algorithms. The error bar has also been shown in each panel.

To demonstrate PEX is an outstanding approach to search in the local landscape around the wild-type sequence, we consider two simple alternative methods to restrict the exploration scope:

#### 1. Reject by Distance (RD)

Rejecting sequences with more than 10 mutations since Figure 5 has shown that 10 mutation sites are sufficient to cover the high-fitness sequences found by PEX. This simple heuristics is previously used by Biswas et al. (2021) where the distance threshold is named by the “trust radius” of the wild type sequence.

#### 2. Lower Confidence Bound (LCB)

Using lower confidence bound that minuses the standard deviation of the ensemble predictions from the average to make the exploration algorithm be conservative on landscape regions with high model uncertainty. Similar approaches are used in the literature of model-based reinforcement learning (Kidambi et al., 2020; Yu et al., 2020).

The experiment results are presented in Table 3. These simple tricks work well in a few cases but cannot fundamentally improve the performance as what is achieved by PEX.

**Table 3.** Comparison to two simple alternative methods to restrict the exploration scope. The learning curves of all eight landscapes are included in Appendix E.

| Algorithm | avGFP | AAV | TEM | E4B |
| --- | --- | --- | --- | --- |
| PEX | **2.97** | **2.52** | 0.19 | **2.21** |
| AdaLead + RD | 2.53 | -0.12 | **0.20** | 1.23 |
| AdaLead + LCB | 2.67 | -2.30 | 0.09 | -0.06 |
| AdaLead + RD + LCB | 2.59 | 1.25 | **0.20** | 1.18 |
| CbAS + RD | 2.28 | -1.51 | 0.12 | 0.45 |
| CbAS + LCB | 2.30 | -2.37 | 0.13 | 0.24 |
| CbAS + RD + LCB | 2.27 | -0.60 | 0.15 | 0.36 |

## 6. Conclusion

This paper propose a novel model-guided directed evolution algorithm, called proximal exploration (PEX), that models the protein sequence design problem by finding the proximal maximization point near the wild-type sequence. It derives a regularized objective function to locally search for sequences with low mutation count upon the wild-type sequence. Based on the localized property of PEX, we propose MuFacNet to specialize in the landscape modeling near the wild type, which factorizes the epistasis effects of mutations to the interactions among the mutated amino acids. In experiments, the integration of PEX and MuFacNet shows great performance improvement compared to prior model-guided protein design baselines in terms of both sample efficiency and the quality of designed sequences.

## A. Experiment Setting and Implementation Details

### A.1. Experiment Setting

As discussed in section 5.1, we collect eight large-scale datasets from previous experimental studies of protein landscape and use TAPE (Rao et al., 2019) as an oracle model to simulate the landscape. Since the wild-type sequences in most datasets have already achieved a high fitness score, we relocate the starting sequence of the search algorithm to the sequence with the lowest fitness in the dataset. It means the starting sequences of these benchmarks are not the true wild-type sequences in the natural environment.

The exploration protocol can only interact with the simulated landscape through batch queries and cannot obtain any extra information about the black-box oracle. The exploration procedure contains 10 rounds of black-box queries. Each batch contains 100 sequences. In Figure 4, we plot the cumulative maximum fitness scores achieved by every algorithm, i.e, the score at round *t* is the maximum fitness scores overall sequences queried in round 1 to *t*.

### A.2. Implementation Details of PEX

The query generation procedure of PEX contains several steps at each round:

1. As a special case, the first round of query generation is completed by randomly mutating the wild-type sequence without accessing to the surrogate model, since the landscape exploration algorithm has not accumulated any protein-fitness data from the interaction with the black-box oracle. The main procedure of PEX is launched at the second round of query generation.
2. To reduce the computation burden of the surrogate model, we perform a heuristics to make the size of candidate pool 𝒟^pool^ manageable. We first compute the proximal frontier of the measured sequence buffer 𝒟. *i*.*e*., we compute the upper convex hull of on the coordinate system constructed by (1) the hamming distance *d*(*s, s*^wt^) to the wild-type sequence *s*^wt^ and (2) the fitness score *f*(*s*) obtained from the landscape oracle. We use Andrew’s monotone chain convex hull algorithm (Andrew, 1979) as follows:
  a. First, we sort the sequences in buffer 𝒟in the increasing order of *d*(*s, s*^wt^). For sequences with the same value of *d*(*s, s*^wt^), we only preserve the one with largest fitness score *f* (*s*).
  b. We maintain a stack whose the elements constructs a convex hull. Initially, the stack is set to empty.
  c. Then, we sweep the sorted sequences. For each sequence, we will intend to push it to the top of the stack. Before the push operation, we will continually pop the elements on the top of the stack that would violate the convex condition when pushing the new element. The convex condition is checked by computing the cross product.
  d. The prefix of the final stack with monotonic increasing values of *f*(*s*) refer to the proximal frontier prox(*s*^wt^, *f*).
  The above procedure can be completed in *O*(*n* log *n*) time with *n* = | 𝒟^pool^|. We screen the measured sequences near the proximal frontier and only perform random mutations upon these outstanding sequences to construct the candidate pool 𝒟^pool^, since the mutants based on the known proximal frontier are more likely to be selected in the further steps of PEX algorithm.
3. Following line 4-7 of Algorithm 2, we iteratively compute the proximal frontier of the candidate pool, add the sequences lying on the frontier to the query batch, and remove them from the candidate pool. This procedure is repeated until we gather sufficient sequences in the query batch. The proximal frontier is also computed by Andrew’s monotone chain convex hull algorithm.

### A.3. Architecture of MuFacNet

The input of MuFacNet is a set of context window of the mutation sites. Each context window is a one-hot encoding for the sequence fragment centered at the corresponding mutation site. The context window size is *L*_*c*_ = 21 (*i*.*e*., radius= 10).

We use Adam optimizer with a learning rate 10^−3^ to train the fitness prediction. The loss function is the mean squared error. We stop network training when the training loss does not decrease in 10 epochs. This training procedure is applied to all models considered in this paper.

**Table 4.** The network architecture of MuFacNet.

|  |  |
| --- | --- |
| **Input:** | $K$ fragments with shape $L_c \times 20$ |
| 1D-CNN | number of filters: 32, kernel size: 5<br>ReLU activation |
| 1D-CNN | number of filters: 32, kernel size: 5<br>ReLU activation |
|  | Global Max-Pooling |
| Dense | output size: 128<br>ReLU activation |
| Dense | output size: 32 |
| **Local Features:** | $K$ feature vectors with length 32 |
|  | Sum-Pooling |
| **Joint Features:** | a feature vector with length 32 |
| Dense | output size: 128<br>ReLU activation |
| Dense | output size: 128<br>ReLU activation |
| Dense | output size: 1 |
| **Fitness Prediction:** | a scalar value |

## B. Experiments based on ESM-1b

We consider an alternative oracle model built on the features produced by ESM-1b (Rives et al., 2021). We adopt the ESM-1b feature with dimension 1280 and train an Attention1D decoder to predict the fitness value.

**Figure 6.**
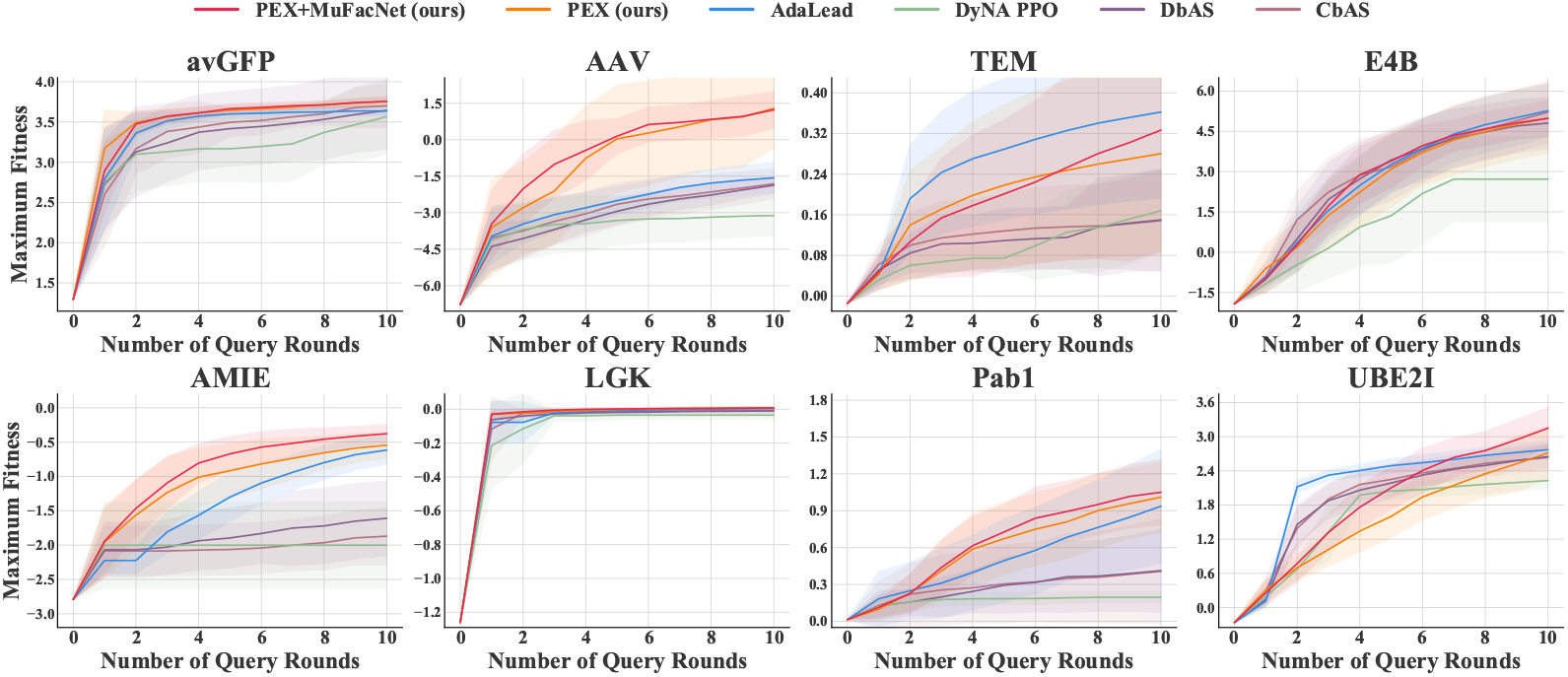
Learning curves of PEX and baselines on a suite of protein engineering tasks simulated by oracle models based on ESM-1b.

## C. Learning Curves of the Experiments in Table 1

**Figure 7.**
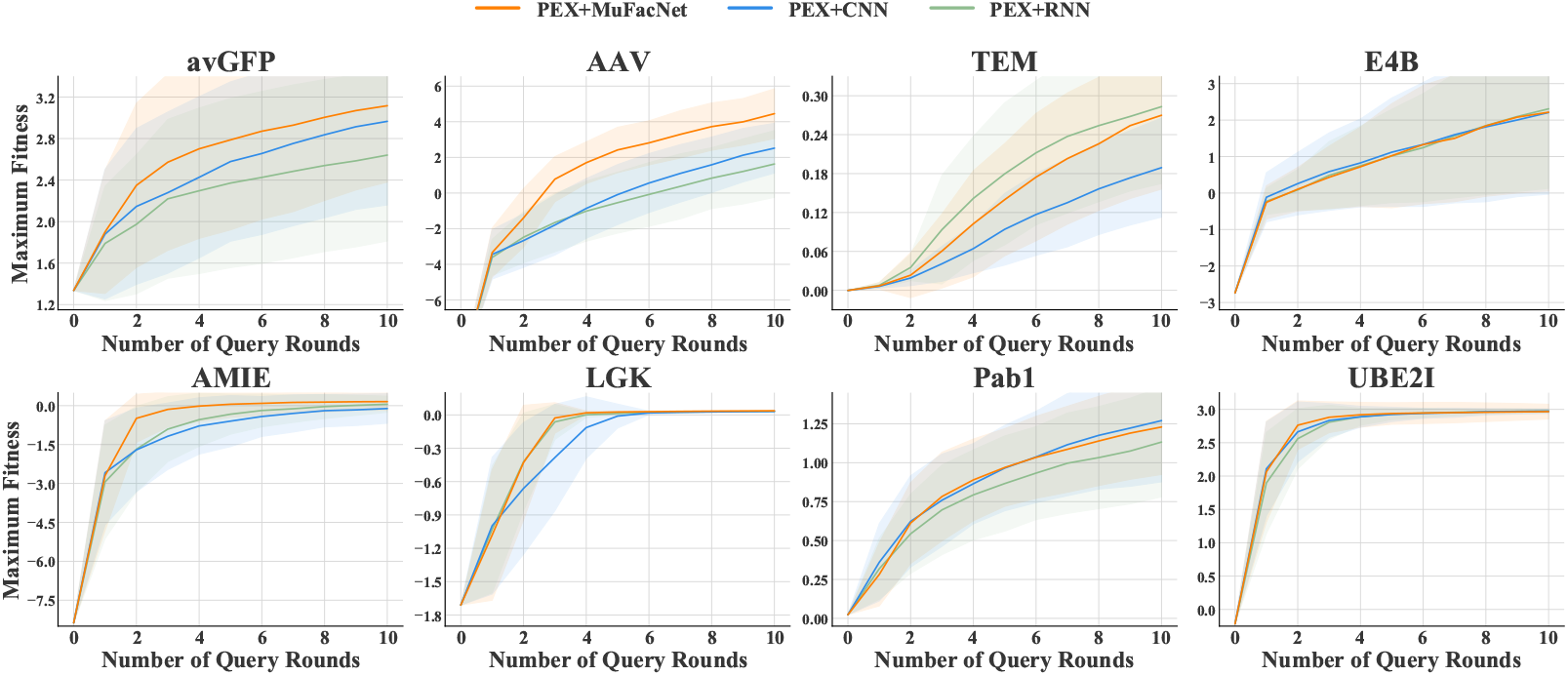
Learning curves of PEX with MuFacNet, CNN, and RNN.

**Figure 8.**
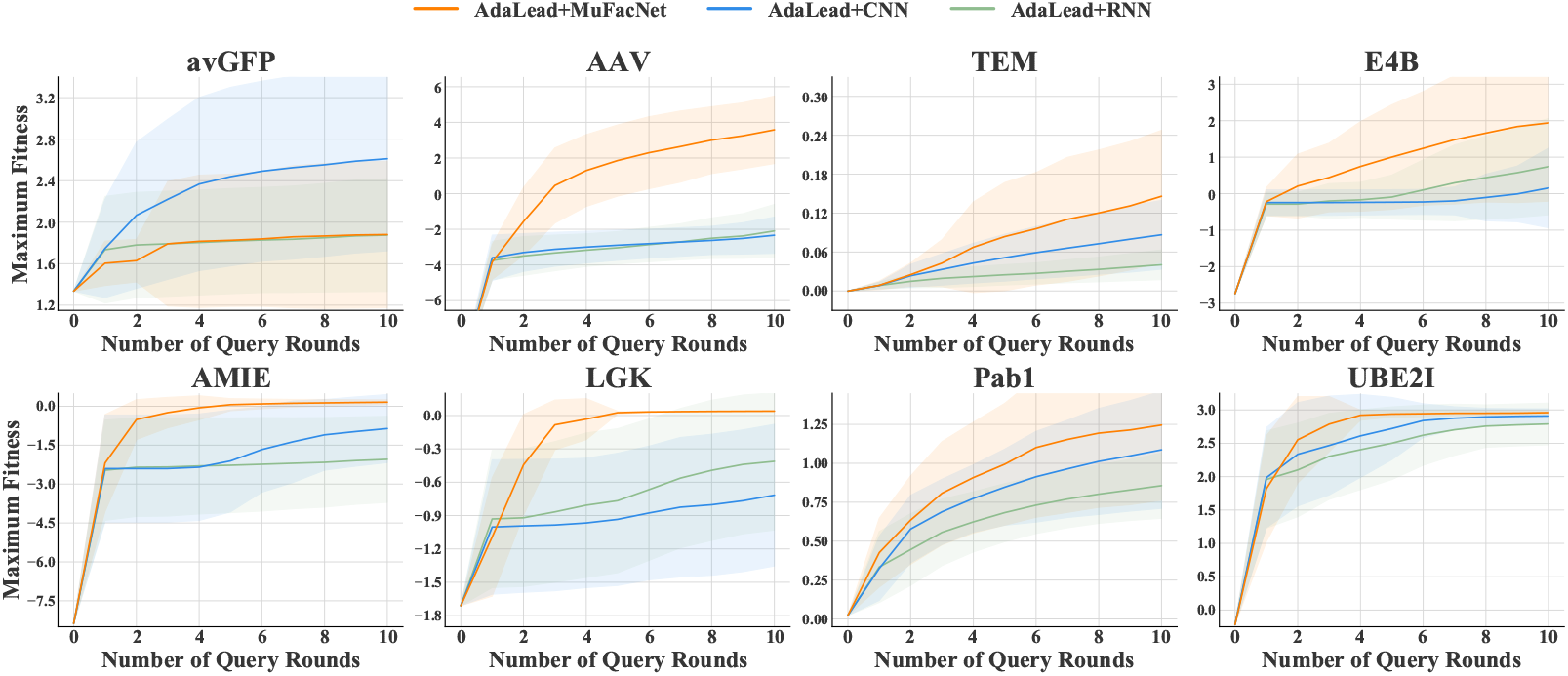
Learning curves of AdaLead with MuFacNet, CNN, and RNN.

**Figure 9.**
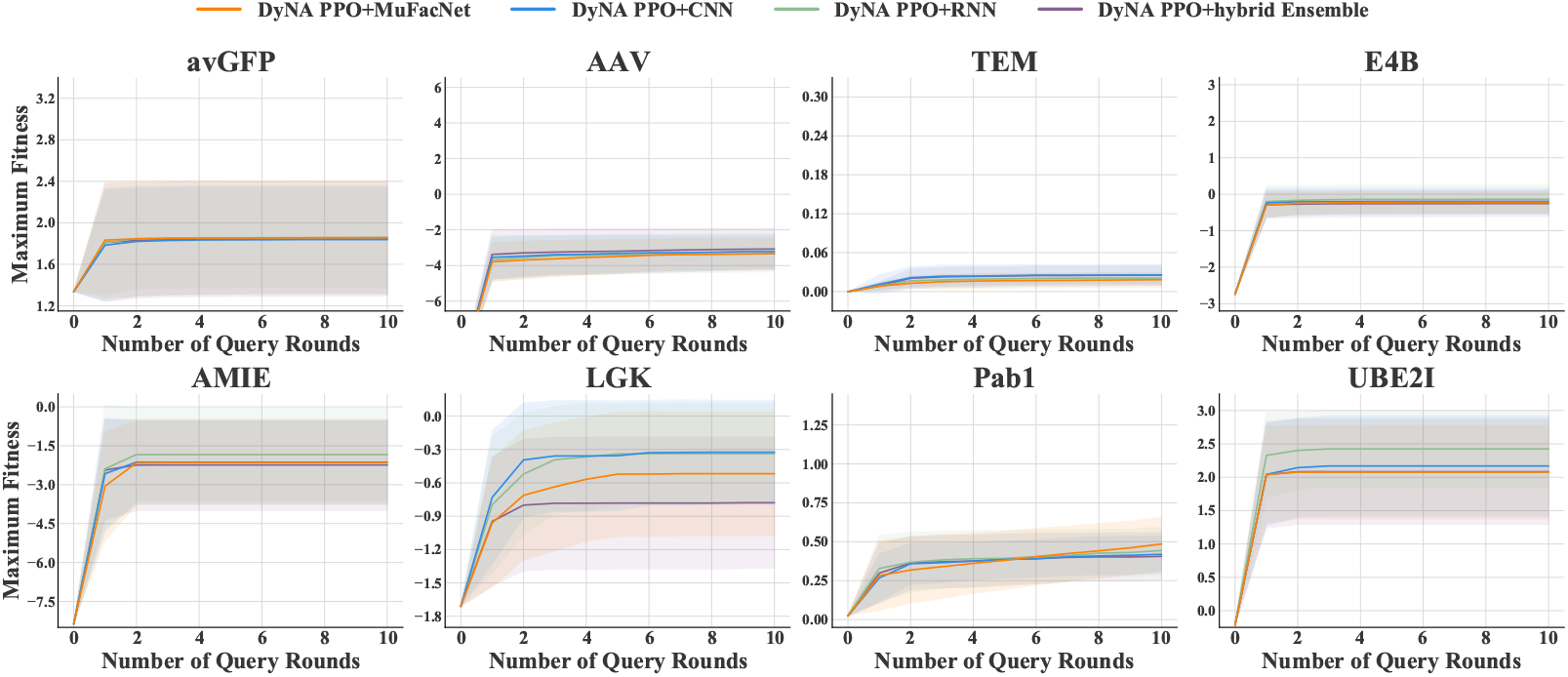
Learning curves of DyNA PPO with MuFacNet, CNN, RNN, and its default hybrid ensemble.

**Figure 10.**
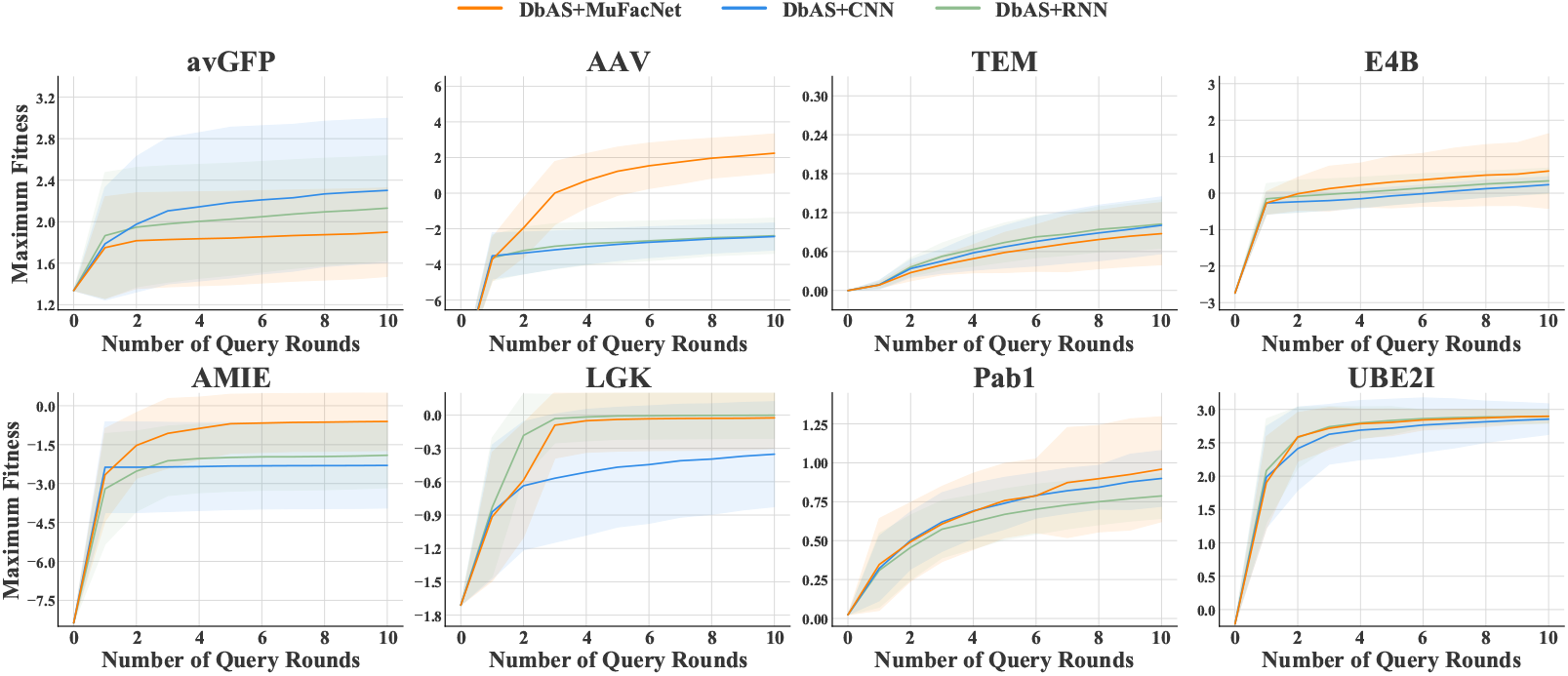
Learning curves of DbAS with MuFacNet, CNN, and RNN.

**Figure 11.**
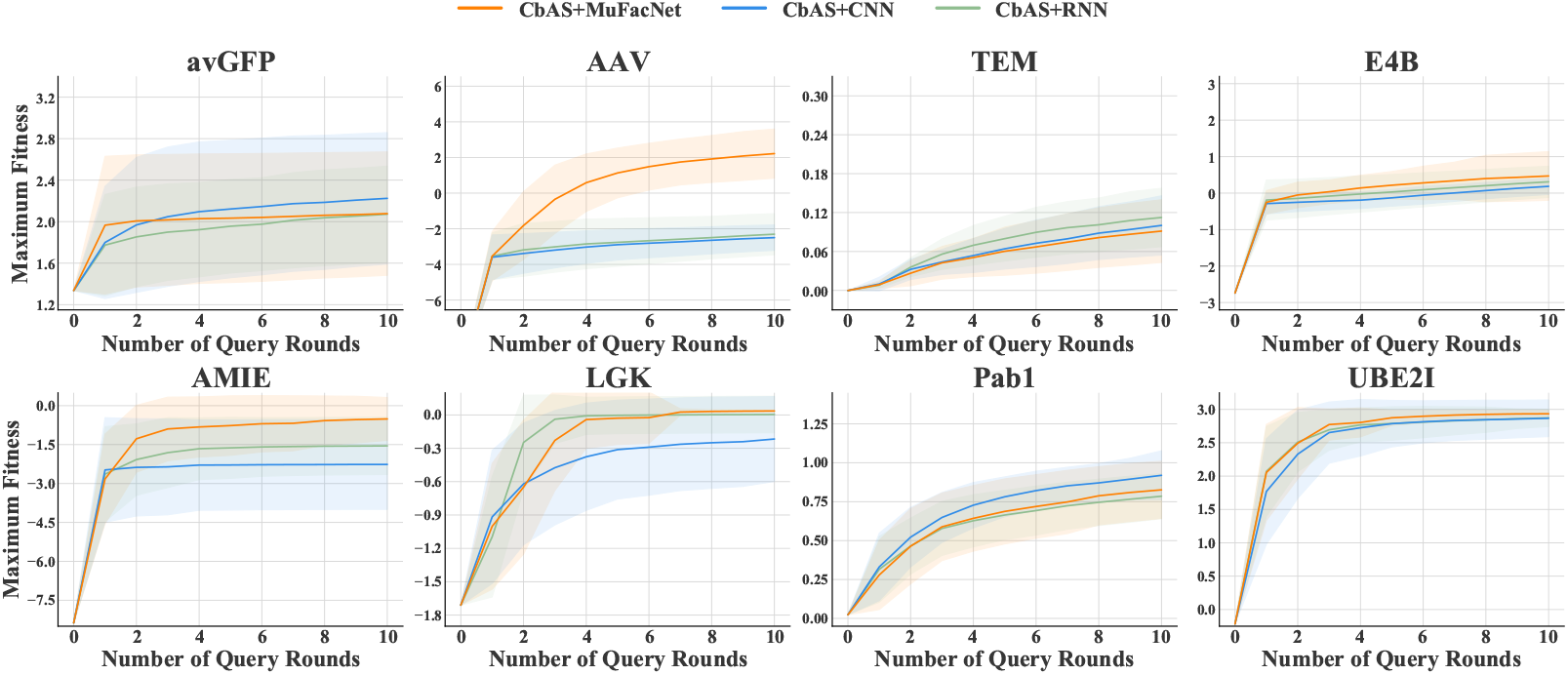
Learning curves of CbAS with MuFacNet, CNN, and RNN.

## D. Learning Curves of the Experiments in Table 2

**Figure 12.**
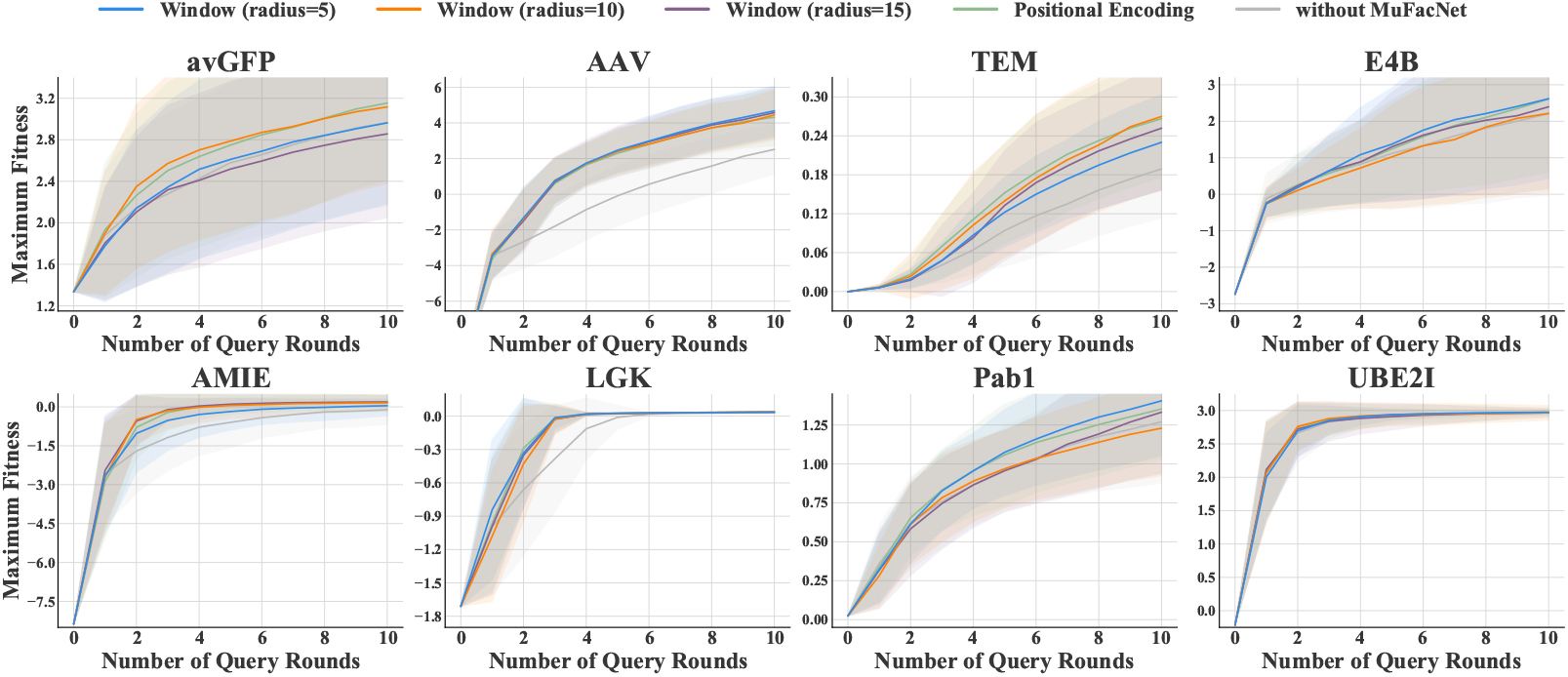
Learning curves of ablation studies on the representation of the context information of each single amino-acid mutation site.

## E. Learning Curves of the Experiments in Table 3

**Figure 13.**
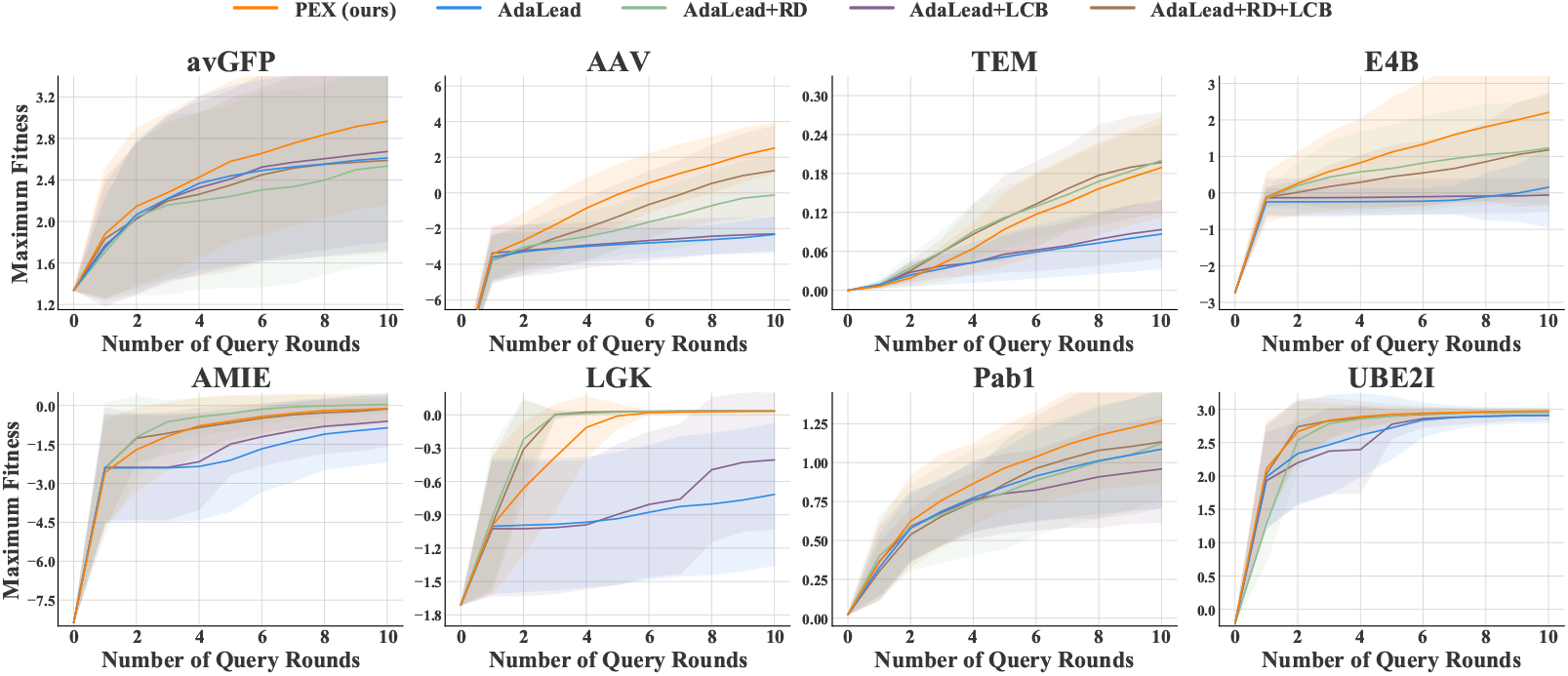
Learning curves of AdaLead with two simple methods to restrict the exploration scope.

**Figure 14.**
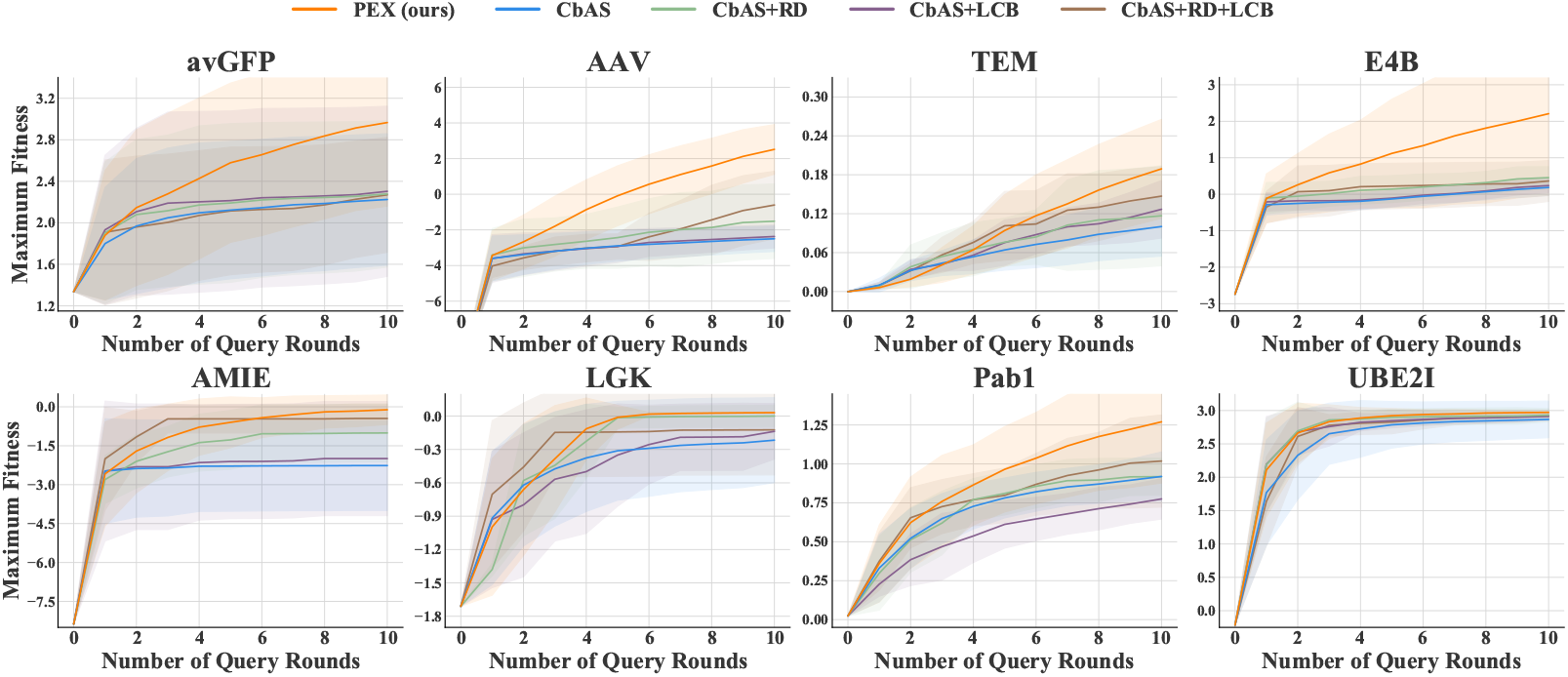
Learning curves of CbAS with two simple methods to restrict the exploration scope.

